# ACTION AFFORDANCE AFFECTS PROXIMAL AND DISTAL GOAL-ORIENTED PLANNING

**DOI:** 10.1101/2021.07.27.454022

**Authors:** Ashima Keshava, Nina Gottschewsky, Stefan Balle, Farbod Nosrat Nezami, Thomas Schüler, Peter König

**Affiliations:** University of Osnabrück, Germany; Halocline GmbH & Co. KG, Germany; University of Osnabrück University Medical Center, Hamburg-Eppendorf, Germany

**Keywords:** anticipatory behavior, action-oriented attention, active inference, eye-tracking, virtual reality, natural cognition

## Abstract

Seminal studies on human cognitive behavior have been conducted in controlled laboratory settings, demonstrating that visual attention is mainly goal-directed and allocated based on the action performed. However, it is unclear how far these results generalize to cognition in more naturalistic settings. The present study investigates active inference processes revealed by eye movements during interaction with familiar and novel tools with two levels of realism of the action affordance. We presented participants with 3D tool models that were either familiar or unfamiliar, oriented congruent or incongruent to their handedness, and asked participants to interact with them by lifting or using. Importantly, we used the same experimental design in two setups. In the first experiment, participants interacted with a VR controller in a low realism environment; in the second, they performed the task with an interaction setup that allowed differentiated hand and finger movements in a high realism environment. We investigated the differences in odds of fixations and their eccentricity towards the tool parts before action initiation. The results show that participants fixate more on the tool’s effector part before action initiation for the use task for unfamiliar tools. Furthermore, with more realistic action affordances, subjects fixate more on the tool’s handle as a function of the tool’s orientation, well before the action was executed. Secondly, the spatial viewing bias on the tool reveals early fixations are influenced by the task and the familiarity of the tools. In contrast, later fixations are associated with the manual planning of the interaction. In sum, the findings from the experiments suggest that fixations are made in a task-oriented way to plan the intended action well before action initiation. Further, with more realistic action affordances, fixations are made towards the proximal goal of optimally planning the grasp even though the perceived action on the tools is identical for both experimental setups. Taken together, proximal and distal goal-oriented planning is contextualized to the realism of action/interaction afforded by an environment.

## 1 Introduction

A longstanding goal of the cognitive sciences is to understand cognition, behavior, and experience as it unfolds in the natural world (Parada and Rossi, 2020). Given the technological advancements made in the last decade, there are few methodological roadblocks to understanding natural cognition where laboratory studies can be extended to naturalistic settings and hopefully lead towards new insights (Ladouce et al., 2016; Parada, 2018). More recently, a pragmatic turn has emerged in the field where there is a greater push towards incorporating the body and bodily actions to infer cognitive function (Engel et al., 2013).

Human tool use is an explicitly natural cognitive function that involves the transfer of proximal goals (e.g., placement of grasp) to distal goals for the tool (Arbib et al., 2009). Moreover, simple tools fundamentally expand the body representations to include representations of the tool in the peripersonal space (Berti and Frassinetti, 2000; Farnè et al., 2005; Maravita et al., 2002). Furthermore, tool use is differentiated from other object-based actions where the tool is “acted with” (Johnson and Grafton, 2003) and requires semantic knowledge of the tool as well as the necessary skill to perform actions with it (Johnson-Frey, 2004). Hence, tool use involves complex behaviors ranging from cognitive and semantic reasoning to perceptual and motor processing.

When using tools, a wealth of information is parsed to produce the relevant action. The semantic knowledge associated with the tool helps understand how it is used, the mechanical knowledge maps the physical properties of the tool for potential usage, and finally, sensorimotor knowledge helps decipher possible movements required to use the tool (Baumard et al., 2014). The amalgamation of these knowledge sources (which can be unique to a tool) necessitates planning any action associated with the tool. When this knowledge is not readily available, inferential processes must be deployed to deduce the relevant action.

In naturalistic settings, studies have shown that eye movements are made to locations in the scene in anticipation of the following action (Hayhoe, 2004; Land and Hayhoe, 2001; Pelz and Canosa, 2001). Belardinelli et al. (2016b) showed that eye movements are goal-oriented and are modulated in anticipation of the object interaction task. There is strong evidence that task plays a vital role in how the eyes scan the scene and are differentiated between passive viewing and pantomimed interaction (Belardinelli et al., 2015). Similarly, Keshava et al. (2020) showed that rudimentary object interactions can be decoded using eye-movement data alone. rudimentary object interactions can be decoded using eye-movement data alone. Even in the absence of an interaction, task relevance plays an important role (Castelhano et al., 2009; Henderson and Hayes, 2017). These studies point towards gaze control being the consequence of knowledge and task-driven predictions (Henderson, 2017).

Moreover, Belardinelli et al. (2016a) investigated the role of anticipatory eye movements when interacting with familiar and unfamiliar tools in a controlled lab setting. These tools had differentiable parts: tool handle and effector. The results showed that in the case of unfamiliar tools, preparatory eye movements are made to the tool-effector to extract the mechanical properties of the tool as the semantic information was not readily available. This effect was enhanced when subjects were asked to perform tool-specific movements instead of a generic action of lifting the tool by the handle. The authors, hence, concluded that eye movements are used to actively infer the appropriate usage of the tools from their mechanical properties. In the study, the tools were presented as images on a screen, and participants pantomimed lifting or using the tool. While the study revealed valuable insights into anticipatory gaze control, a question remains if these results are part of natural cognition and can be reproduced in more realistic environments.

Herbort and Butz (2011) further investigated the interaction of habitual and goal-directed processes that affect grasp selection while interacting with everyday objects. They presented objects in different orientations and showed that grasp selection depended on the overarching goal of the movement sequence dependent on the object’s orientation. Belardinelli et al. (2016b) further showed that fixations have an anticipatory preference for the region where the index finger is placed. Consequently, the location of fixations is predictive of both proximal goals of manual planning and task-related distal goals.

When studying anticipatory behaviors corresponding to an action, one must also ask whether symbolized action is enough and how real the action should be. Króliczak et al. (2007) showed brain areas typically involved in real actions are not driven by pantomimed actions and that pantomimed grasps do not activate the object-related regions within the ventral stream. Similarly, Hermsdörfer et al. (2012) showed a weak correlation between the hand trajectories for pantomimed and actual tool interaction. These studies indicate that the realism of sensory and tactile feedback while acting (e.g., a grasp) can be an essential factor when studying anticipatory behavioral control.

In virtual reality (VR), realistic actions can be studied by simulating an interaction within an environment. Using interfaces such as VR controllers, ego-centric visual feedback of a hand can be simulated. These interfaces usually consist of hand-held devices that are tracked in space and through which different actions are controlled by pressing buttons. One advantage of controller-based VR interaction is the possibility of haptic feedback. One disadvantage is that the hand posture while holding the controller does not always correspond to the user’s virtual visual feedback when the simulated hand performs the action. Conversely, camera-based interaction interfaces such as LeapMotion, capture the real-time movements of the user’s hand and use finger gestures, like wrap grasp or pinch grasp, to control different actions in the environment. These interfaces give the user realistic visual feedback of their finer hand and finger movements, while they can not give haptic feedback. Consequently, the chosen method of interaction in VR can afford different levels of realism and could elicit different behavioral responses.

In the present study, we investigated anticipatory gaze control in two different experiments. We were interested in the extent to which the realism of the action affordance and the environment modified the results shown by Belardinelli et al. (2016a). We asked participants to lift or use 3D models of tools in VR that were categorized as familiar or unfamiliar. Additionally, we extended the experimental design to include the tool handle’s spatial orientation, congruent or incongruent to the subjects’ handedness.

In experiment-I, subjects performed the experiment in a low realism environemnt and action affordance and interacted with the tool models using a VR controller, which mimicked grasp in the virtual environment by pulling the index finger. In experiment-II, subjects were immersed in a high realism setting where they interacted with the tools using LeapMotion, which required natural hand and finger movements. Thus, the action affordance appeared closer to the real world. With this experimental design, we investigate the influence of task, tool familiarity, the spatial orientation of the tool, and, notably, the impact of the realism of the action affordance.

## 2 Methods

### 2.1 Experimental Task

Subjects were seated in a virtual environment where they had to interact with the presented tool by either lifting or pretending its use. The time course of the trials is illustrated in Figure 1A. At the start of a trial, subjects saw the cued task for 2 sec after which the cue disappeared, and a tool appeared on the virtual table. Subjects were given 3 sec to view the tool, after which there was a beep (go cue) which indicated that they could start manipulating the tool based on the cued task. Subjects were seated in a virtual environment where they had to interact with the presented tool by either lifting or pretending its use. After interacting with the tool, subjects pressed a button on the table to start the next trial.

**Figure 1:**
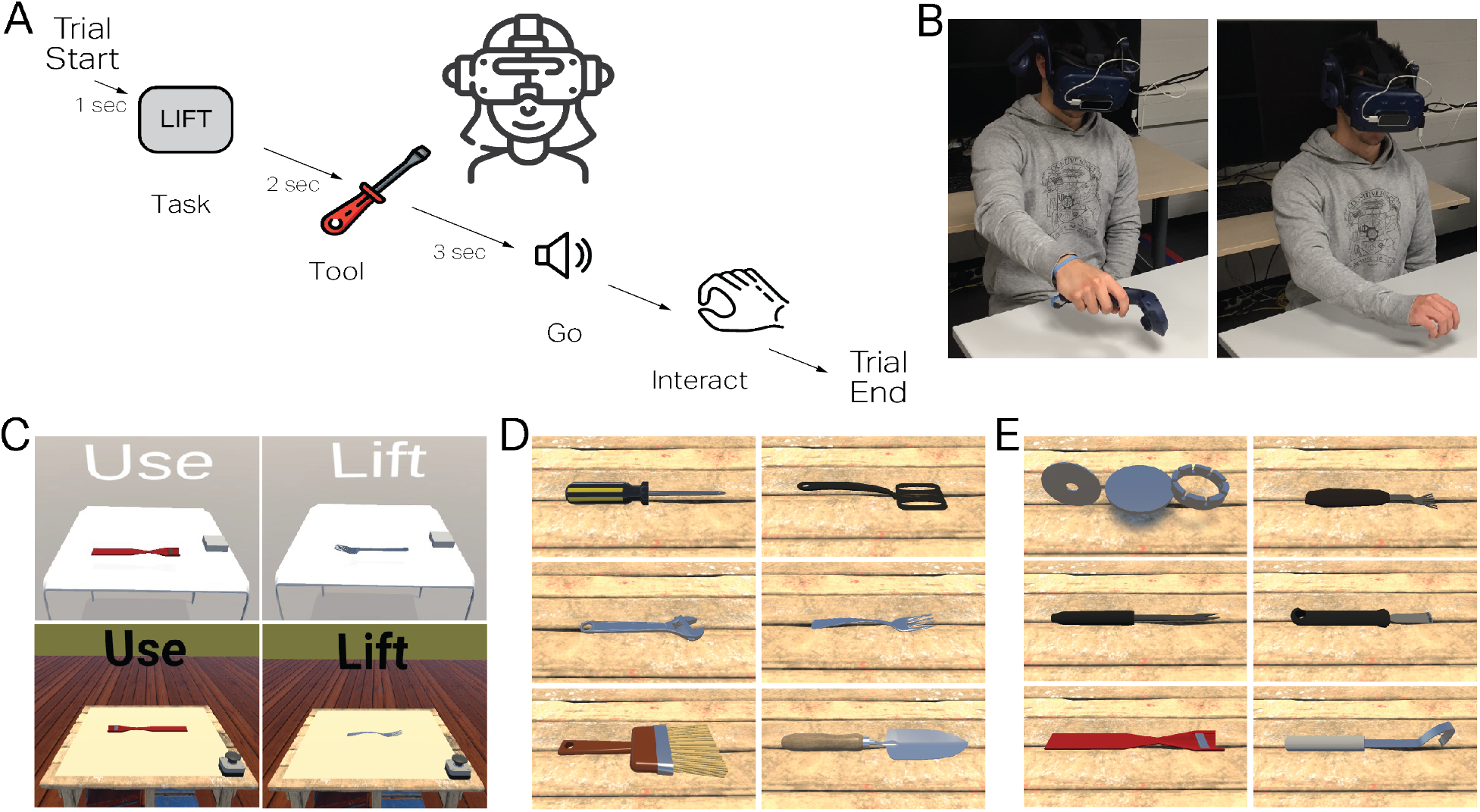
Experimental Task. In two virtual environments participants interacted with tools in two ways (LIFT, USE). The tools were categorized based on familiarity (FAMILIAR, UNFAMILIAR) and presented to the participants in two orientations (HANDLE LEFT, HANDLE RIGHT). The two virtual environments differed based on the mode of interaction and perceived realism, wherein in one experiment, subjects’ hand movements were rendered virtually using the HTC-VIVE controllers. In the other experiment, the hands were rendered using LeapMotion, allowing finer hand and finger movements. **Panel A** shows the timeline of a trial. **Panel B** shows a subject in real-life performing the task in the two experiments. **Panel C** shows the differences in realism in the two experiments; TOP panels correspond to experiment with the controllers, the USE and LIFT conditions for an UNFAMILIAR and FAMILIAR tool, respectively with the tool handles presented in two different orientations. BOTTOM panels illustrate the three different conditions in a more realistic environment with LeapMotion as the interaction method. **Panel D** Familiar tools, from top-left: screwdriver, spatula, wrench, fork, paintbrush, trowel. **Panel E** Unfamiliar tools, from top-left: spoke-wrench, palette knife, daisy grubber, lemon zester, flower cutter, fish scaler.

### 2.2 Participants

For experiment-I with the HTC Vive controller’s interaction method, we recruited 18 participants (14 females, mean age=23.68, SD=4.05 years). For experiment-II with the interaction method of LeapMotion, we recruited 30 participants (14 female, mean age=22.7, SD=2.42 years). All participants were recruited from the University of Osnabrück and the University of Applied Sciences Osnabrück. Participants had a normal or corrected-to-normal vision and no history of neurological or psychological impairments. All of the participants were right-handed. They either received a monetary reward of C10 or one participation credit per hour. Before each experimental session, subjects gave their informed consent in writing. They also filled out a questionnaire regarding their medical history to ascertain they did not suffer from any disorder/impairments which could affect them in the virtual environment. Once we obtained their informed consent, we briefed them on the experimental setup and task.

### 2.3 Experimental Design and Procedure

The two experiments differed based on the realism of the action affordance and the environment. Figure 1B illustrates the physical setup of the participants for the two experiments. In experiment-I, subjects interacted with the tool models using the HTC Vive VR controllers. While in experiment II, subjects’ hand movements were captured by LeapMotion. Figure 1C illustrates two exemplar trials from the experiments. We used a 2×2×2 experimental design for both experiments, with factors task, tool familiarity, and handle orientation. Factor task had two levels: LIFT and USE. In the LIFT conditions, we instructed subjects to lift the tool to their eye level and place it back on the table. In the USE task, they had to pantomime using the tool to the best of their knowledge. Factor familiarity had two levels, FAMILIAR and UNFAMILIAR, which corresponded to tools either being everyday familiar tools or tools that are not seen in everyday contexts and are unfamiliar. The factor handle orientation corresponded to the tool handle, which was presented to the participants either on the LEFT or the RIGHT. Both experiments had 144 trials per participant, with an equal number of trials corresponding to the three factors. Subjects performed the trials over six blocks of 24 trials each. We measured the eye movements and hand movements simultaneously while subjects performed the experiment. We calibrated the eye-trackers at the beginning of each block and ensured that the calibration error was less than 1 degree of the visual angle. At the beginning of the experiment, subjects performed three practice trials with a hammer to familiarize themselves with the experimental setup and the interaction method. Each experiment session lasted for approximately an hour. After that, subjects filled out a questionnaire to indicate their familiarity with the 12 tools used in the experiment. They responded to the questionnaire based on a scale of 5-point Likert-like scale where 1 corresponded to “I have never used it or heard about it,” and 5 referred to “I see it every day or every week.”

### 2.4 Experimental Stimuli

The experimental setup consisted of a virtual table that mimicked the table in the real world. The table’s height, width, and length were 86cm, 80cm, and 80cm, respectively. In experiment-I, subjects were present in a bare room with grey walls and constant illumination. They sat before a light grey table, with a dark grey button on their right side to indicate the end of the trial. Similarly, in experiment-II, subjects were present in a more immersive, realistic room. They sat in front of a wooden workbench with the exact dimensions of the real-world table and a buzzer on the right to indicate the end of a trial. We displayed the task (USE or LIFT) over the desk 2m away from the participants for both experiments. For both experiments, we used the tool models as presented in Belardinelli et al. (2016a). Six of the tools were categorized as familiar (Figure 1D) and the other six as unfamiliar (Figure 1E). We further created bounding box colliders that encapsulated the tools to capture the gaze position on the tool models. The mean length of the bounding box was 34.04cm (SD=5.73), mean breadth=7.60cm (SD=3.68) and mean height= 4.17cm (SD=2.13). To determine the tool effector and tool handle regions of interest, we halved the length bounding box colliders from the center of the tool and took one half as the effector and the other half as the handle. This way we refrained from making arbitrary-sized regions-of-interest for the different tool models.

### 2.5 Apparatus

For both experiments, we used an HTC Vive head-mounted display (HMD)(110° field of view, 90Hz, resolution 1080 x 1200 px per eye) with a built-in Tobii eye-tracker ^1^. The HTC Vive Lighthouse tracking system provided positional and rotational tracking and was calibrated for a 4m x 4m space. For calibration of the gaze parameters, we used 5-point calibration function provided by the manufacturer. To make sure the calibration error was less than 1°, we performed a 5-point validation after each calibration. Due to the nature of the experiments, which allowed a lot of natural head movements, the eye tracker was calibrated repeatedly during the experiment after each block of 36 trials. We designed the experiment using the Unity3D game engine ^2^ (v2019.2.14f1) and controlled the eye-tracking data recording using HTC VIVE Eye Tracking SDK SRanipal^3^ (v1.1.0.1)

For experiment-I, we used HTC Vive controller^4^ (version 2.5) to interact with the tool. The controller in the virtual environment was rendered as a gloved hand. When participants pulled the trigger button of the controller with their right index finger, their right virtual hand made a power grasp action. To interact with the tools, subjects pulled the trigger button of the controller over the virtual tools and the rendered hand grasped the tool handle.

Similarly, in experiment-II, we used LeapMotion^5^ (version 4.4.0) to render the hand in the virtual environment. Here, subjects could see the finer hand and finger movements of their real-world movements rendered in the virtual environment. When participants made a grasping action with their hand over the virtual tool handle, the rendered hand grasped the tool handle in the virtual environment.

### 2.6 Data pre-processing

#### 2.6.1 Gaze Data

As a first step, using eye-in-head 3D gaze direction vector for the cyclopean eye we calculated the gaze angles in degrees for the horizontal *θ*_h_ and vertical *θ*_v_ directions. All of the gaze data was sorted by the timestamps of the collected gaze samples. The 3D gaze normals are represented as (*x, y, z*) a unit vector that defines the direction of the gaze in VR world coordinates. In our setup, the x coordinate corresponds to the left-right direction, y in the up-down direction, z in the forward-backward direction. The formulas used for computing the gaze angles are as follows:

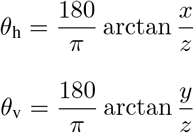

Next, we calculated the angular velocity of the eye in both the horizontal and vertical coordinates by taking a first difference of the angular velocity and dividing by the difference between the timestamp of the samples using the formula below:

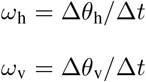

Finally, we calculated the magnitude of the angular velocity (*ω*) at every timestamp from the horizontal and vertical components using:

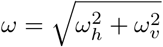

To classify the fixation and saccade-based samples, we used an adaptive threshold method for saccade detection described by Voloh et al. (2020). We selected an initial saccade velocity threshold *θ*_0_ of 200 °/sec. All eye movement samples with an angular velocity of less than *θ*_0_ were used to compute a new threshold *θ*_1_. *θ*_1_ was three times the median absolute deviation of the selected samples. If the difference between *θ*_1_ and *θ*_0_ was less than 1 °/sec *θ*_1_ was selected as the saccade threshold else, *θ*_1_ was used as the new saccade threshold and the above process was repeated. This was done until the difference between *θ*_n_ and *θ*_n+1_ was less than or equal to 1 °/sec. This way we arrived at the cluster of samples that belonged to fixations and the rest were classified as saccades.

After this, we calculated the duration of the fixations and removed those fixations that had a duration less than 50 ms or were larger than 3.5 times the median absolute deviation of the fixation duration. For further data analysis, we only considered those fixations that were positioned on the 3D tool models. We further categorized the fixations based on their position on the tool, i.e., whether they were located on the effector or handle of the tool.

### 2.7 Data Analysis

#### 2.7.1 Odds of Fixations in favor of tool effector

After cleaning the dataset, we were left with 2174 trials from 18 subjects in experiment-I and 3633 trials from 30 subjects in experiment-II. For both experiments, we analysed the fixations in the 3 second period from the tool presentation till the go cue. For the two experiments, we modeled the linear relationship of the log of odds of fixations on the effector of the tools and the task cue (LIFT, USE), the familiarity of the tool (FAMILIAR, UNFAMILIAR), and orientation of the handle (LEFT, RIGHT). All within-subject effects were also modeled with random intercepts and slopes based on the subjects. We were also interested in modeling the random effects based on the tool to assess the differential effects on the individual tools. We did not have enough data to estimate random item effects, so we fitted a random intercept for the 12 tools.

We used effect coding (Schad et al., 2018) to construct the design matrix for the linear model, where we coded the categorical variables LIFT, FAMILIAR, RIGHT to −0.5 and USE, UNFAMILIAR, LEFT to 0.5. This way, we could directly interpret the regression coefficients as main effects. The model fit was performed using restricted maximum likelihood (REML) estimation (Corbeil and Searle, 1976) using the lme4 package (v1.1-26) in R 3.6.1. We used the L-BFGS-B optimizer to find the best fit using 10000 iterations. Using the Satterthwaite method (Luke, 2017), we approximated degrees of freedom of the fixed effects. For both experiments, the Wilkinson notation (Wilkinson and Rogers, 1973) of the model was:

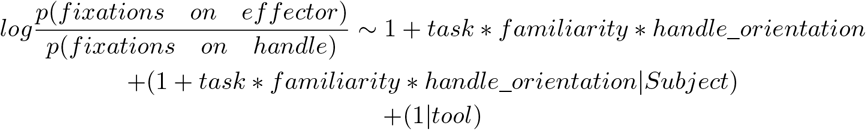

As we used effects coding, we can directly compare the regression coefficients of the two models. The fixed-effect regression coefficients of the two models would describe the differences in log-odds of fixations in favor of tool effector for the categorical variables task, familiarity, and handle orientation.

#### 2.7.2 Spatial bias of fixations on the tools

In this analysis, we wanted to assess the effects of task, tool familiarity, and handle orientation on the eccentricity of fixations on the tools. To do this, we studied the fixations from the time when the tool was visible on the table (3s from the start of trial) till the go cue indicated when subjects could start manipulating the tool. We divided this 3s period into 20 equal bins of 150ms each. For each trial and time bin, we calculated the median distance of the fixations from the tool center. Next, we normalized the distance with the length of the tool so that we could compare the fixation eccentricity across different tools.

To find the time-points where there were significant differences for the 3 conditions and their interactions, we used the cluster permutation method. Here, we use the t-statistic as a test statistic for each time-bin, where t is defined as:

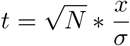

and, x is the mean difference between conditions, and *σ* is the standard deviation of the mean and N is the number of subjects. We used a threshold for t at 2.14 which corresponds to the t-value at which the p-value is 0.05 in the t-distribution. We first found the time-bins where the t-value was greater than the threshold. Then, we computed the sum of the t-values for these clustered time-bins which gave a single value that represented the mass of the cluster. Next, to assess the significance of the cluster, we permuted all the time-bins across trials and subjects and computed the t-values and cluster mass for 1000 different permutations. This gave us the null distribution over which we compared the cluster mass shown by the real data. We considered the significant clusters to have a p-value less than 0.05. In the results, we report the range of the significant time-bins for the 3 different conditions and their interactions and the corresponding p-values.

## 3 Results

The present study investigated the differences in gaze-based strategies dependent on task, tool familiarity, and handle orientation. Here, we investigated two anticipatory gaze-based strategies 3 seconds before action initiation; the odds of fixations in favor of the tool effector and the eccentricity of the fixations through time towards the tool effector. We further compared the differences in two experiments that had the same experimental design but differed in the realism of the action affordance and environment.

First, we were interested in how the participants subjectively assessed the familiarity of the 12 tools. 2A shows the subjective familiarity ratings for each of the familiar and unfamiliar tools used in the study. The mean familiarity rating for familiar tools in experiment-I was 4.55 (SD=0.60) and for unfamiliar tools 1.81 (SD=1.17). In experiment-II, the mean familiarity rating for familiar tools was 4.48 (SD=0.52) and for unfamiliar tools 1.56 (SD=1.04). To determine the differences in the subjective familiarity ratings for the two experiments and our categorization of familiarity, we performed a mixed-ANOVA with familiarity as a within-subject factor and the experiment group as the between-subject factor. We found no differences in the familiarity ratings between the two experiments (F(44)=3.08, p-value=0.08). Furthermore, there were significant differences in the subjective rating of the tools (F(44)=3094.05, p-value<0.001). There were also no significant interactions between the two factors (F(44)=2.52, p-value=0.11). Figure In sum, our experimental condition of familiarity was consistent with the participants’ subjective rating as well.

Next, we wanted to make sure that the differences in the virtual environments did not affect the way subjects allocated their attention to the experimental task. We calculated the mean percentage of fixations positioned on the tool vs. anywhere else in the environment for each subject across trials. Figure 2B shows the percentage of fixations allocated to the tools vs. the environment for the two experiments. For experiment-I with the interaction method of VR controller and a less realistic environment, the mean percentage of fixations on the environment was 0.29 (SD=0.08) and on the tools 0.78 (SD=0.10). Conversely, in experiment-II with LeapMotion as the interaction method and a more realistic environment, the percentage of fixations allocated to the environment was 0.30 (SD=0.15) and on the tools 0.80 (SD=0.14). To test if these differences were significant, we performed a mixed-ANOVA with fixation location as a within-subject factor, the two experiments as a between-subject factor, and the percentage of fixations as the dependent variable. We found no differences in the percentage of fixations between the two experiments (F(47)=2.86, p-value=0.09). There were significant differences in the percentage of fixations located on the tool vs. the environment (F(47)=217.47, p-value<0.001). We did not find any interactions between the two factors (F(47)=0.02, p-value=0.87). These results show that the allocation of attention was primarily task-oriented and was not affected by the differences in the virtual environment of the two experiments.

**Figure 2:**
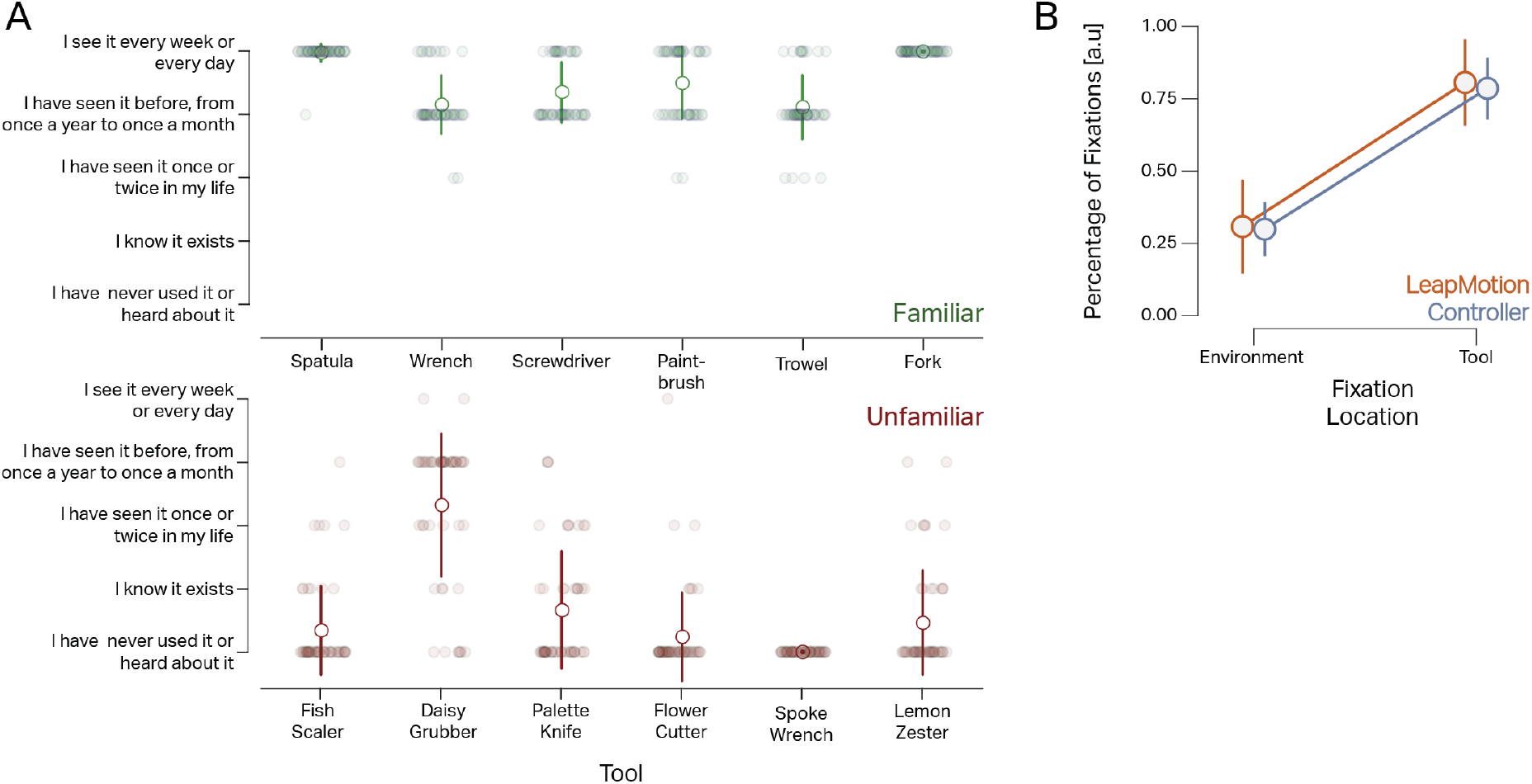
**A)** Participants’ familiarity rating of the tools. Participants provided their subjective rating of familiarity with the 12 tool stimuli on a 5-point Likert scale. The small circles correspond to ratings from individual subjects. The larger circles correspond to the mean rating for each tool, and error bars represent the standard deviation across subjects. **B)** Percentage of fixations allocated to the environment vs. the tools for the two different experiments. The circles correspond to the mean percentage of fixations across subjects, and the error bars represent the standard deviation. As seen here, the realism of the environment did not affect how participants allocated their attention in the experiments.

Next, we compared the log-odds of fixations in favor of the tool effector across the three conditions: task, tool familiarity, and handle orientation in the 3s period when the subjects studied the tool. Figure 3A shows the log-odds of the fixations on the tool effector for experiment-I (with HTC VIVE Controllers) and experiment-II (with LeapMotion).

**Figure 3:**
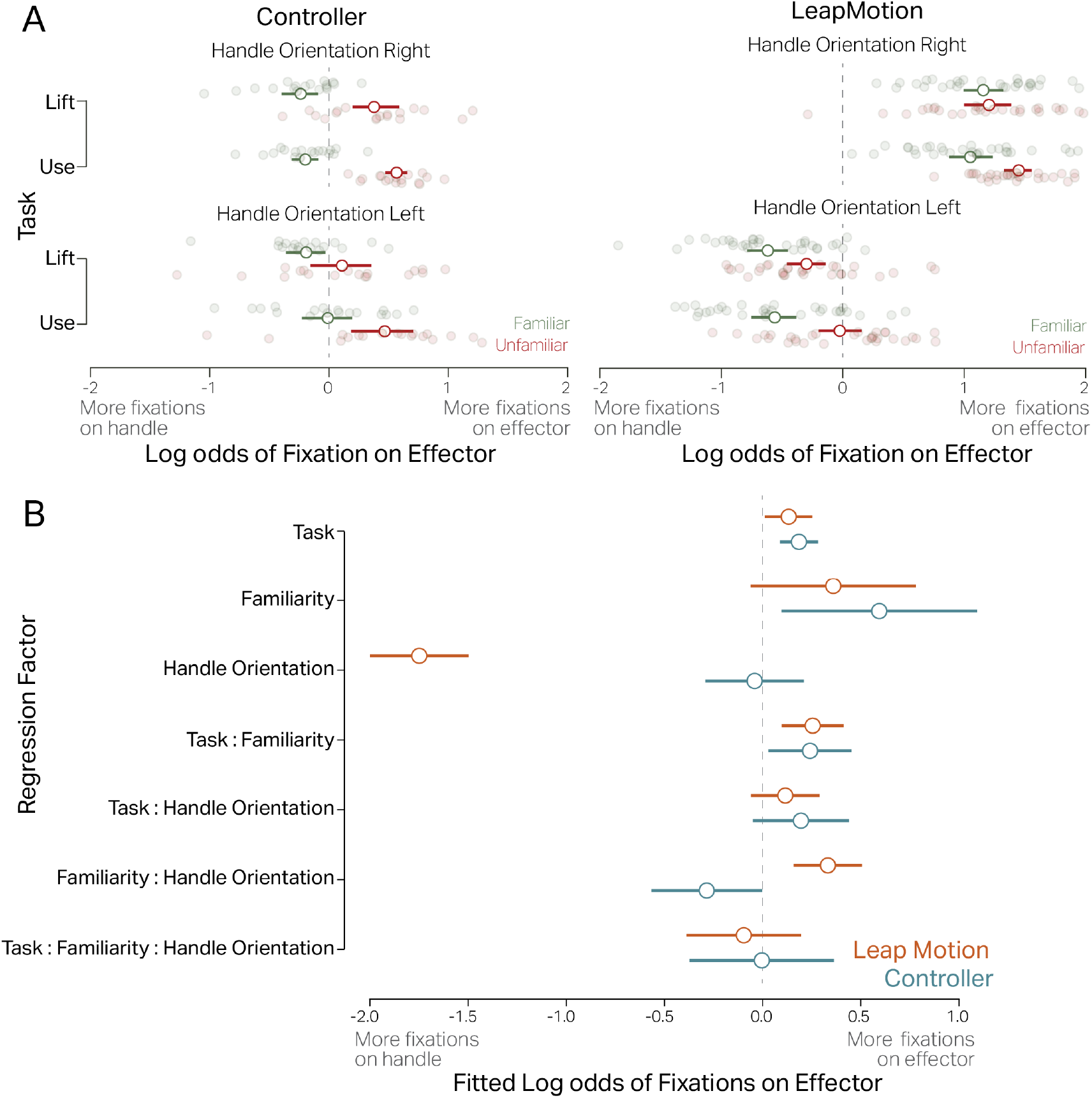
Experiment results. **A) top-left:** shows the log-odds of fixation on effector vs. handle in the controller study when the tool handle is oriented to the right. The log odds on fixations are higher on the effector for unfamiliar tools (red) than the familiar tools (green) for both the LIFT and the USE tasks. **Bottom-left:** log odds of fixation on effector when the tool handle is oriented to the left and is incongruent to the subjects’ handedness. The plot shows that the orientation of the tool does not significantly affect the log-odds fixation on the effector. **Top-right:** the log-odds of fixation on effector in the LeapMotion study when the tool handle is oriented to the right. The log odds of fixations on the effector are higher for unfamiliar tools (red) than the familiar tools (green) and the USE task. **Bottom-right:** log odds of fixation on effector when the tool handle is oriented to the left and is incongruent to the subjects’ handedness. The plot shows that the orientation of the tool results in significant log-odds of fixations over the handle in the LIFT task, while in the USE task and with unfamiliar tools (red) significantly more fixations were on the effector. **B)** The linear regression coefficients for the two experiments. The effect of the task is significant for both experiments with higher log-odds of fixations on the effector. For the factor orientation, the log-odds are significant in the LeapMotion experiment and not for the controller experiment. Similarly, the interaction between task and familiarity is significant for both experiments.

In experiment-I (Figure 3A, left panel), subjects showed a mean log odds of 0.01 (95%CI = [-0.04, 0.08]) for the LIFT task and for the USE task the mean log-odds were 0.19 (95%CI = [0.11, 0.28]). For the FAMILIAR tools, the mean log-odds in favor of the tool effector were −0.16 (95%CI = [−0.23, −0.09]) and for UNFAMILIAR 0.35 (95%CI = [0.25, 0.45]). For the RIGHT oriented tool handle, the mean log-odds were 0.14 (95%CI = [0.06, 0.21]) and for the LEFT oriented tool handle, the mean log-odds were 0.08 (95%CI = [-0.09, 0.26]). To assess the significance of the factors, we used linear mixed models. For the linear model, we used effect coding so the regression coefficients can be directly interpreted as main effects. There was a significant main effect of factor task (USE - LIFT) *β* = 0.18 (95%CI = [0.08, 0.27], t(70.09)=3.7), with a p-value < 0.001. There was a significant main effect of familiarity (UNFAMILIAR - FAMILIAR) *β* = 0.58 (95%CI = [0.09, 1.08], t(10.79)=2.33), with p-value = 0.04. The main effect of handle orientation was not significant (LEFT - RIGHT) *β* = −0.04, (95%CI = [-0.29, 0.21], t(16.94)=-0.32), p-value = 0.75. We found a significant interaction of task and familiarity with *β* = 0.24 (95%CI = [0.03, 0.45], t(25.88)=2.21), p-value = 0.036. The interaction of task and handle orientation was not significant, *β* = 0.19 (95%CI = [-0.05, 0.43], t(21.26)=1.55), p-value = 0.13. The interaction of familiarity and orientation was not significant, *β* = −0.28 (95%CI = [−0.56, −0.004], t(17.07)=-1.99), p-value = 0.06. The 3-way interaction was also not significant, *β* = −0.005 (95%CI = [-0.37, 0.36], t(1695)=-0.02), p-value=0.97.

In experiment-II (Figure 3A, right panel), subjects showed a mean log odds of 0.08 (95%CI = [-0.04, 0.22]) of fixations on the tool effector for the LIFT task and for the USE task the mean log-odds were 0.22 (95%CI = [0.09, 0.36]). For the FAMILIAR tools, the mean log-odds in favor of the tool effector were 0.04 (95%CI = [-0.08, 0.16]) and for UNFAMILIAR 0.25 (95%CI = [0.12, 0.39]). For the RIGHT oriented tool handle, the mean log-odds were 1.18 (95%CI = [1.06, 1.29]) and for the LEFT oriented tool handle, the mean log-odds were −0.32 (95%CI = [−0.45, −0.20]). Using the linear mixed model, we assessed the significance of the three factors. There was a significant main effect of factor task (USE - LIFT) *β* = 0.13 (95%CI = [0.01, 0.25], t(27.28)=2.13) with a p-value = 0.04. The main effect of familiarity (UNFAMILIAR - FAMILIAR) was not significant *β* = 0.35 (95%CI = [-0.06, 0.77], t(10.02)=1.67) with p-value = 0.12. The main effect of handle orientation (LEFT - RIGHT) was significant, *β* = −1.74 (95%CI = [-1.99, −1.48], t(28)=-13.65), p-value <0.001. We found a significant interaction of task and familiarity with *β* = 0.25 (95%CI = [0.09, 0.41], t(43.64)=3.13), p-value = 0.003. The interaction of task and handle orientation was not significant, *β* = 0.11 (95%CI = [-0.06, 0.28], t(31.90)=1.29), p-value = 0.20. Similarly, the interaction of familiarity and orientation was significant, *β* = 0.33 (95%CI = [0.15, 0.50]), p-value = 0.001. The 3-way interaction was also not significant, *β* = −0.09 (95%CI = [-0.39, 0.19], t(2201)=-0.65), p-value=0.51.

Figure 3B summarizes the regression coefficients of the linear model from both experiments. Importantly, we see that the main effect of the task is significant for both experiments. Similarly, the interaction of task and familiarity is significant for both experiments. However, the effect of handle orientation is only significant in experiment-II with the LeapMotion interaction method.

Next, we were interested in the effect of task, tool familiarity, and handle orientation on the eccentricity of the fixations on the tool before action initiation. We calculated the relative distance of fixations from the center of the tool in the 3s period when the subjects studied the tool. We used cluster permutation tests to evaluate the time periods when the effects of the different conditions were significant. As shown in Figure 4A, in experiment-I, the differences in task (USE - LIFT) were significant from 1.05s to 1.95s period, p-value<0.001. Differences in tool familiarity (FAMILIAR - UNFAMILIAR) were significant from 0.15s to 3s with a p-value <0.001. Moreover, the differences in the two orientations (LEFT - RIGHT) were not significant. The interaction of task and familiarity were significant from 0.3s to 2.55s, p-value=0.006. The interactions of task and handle orientation, and the interaction of handle orientation and tool familiarity were not significant in any time period.

**Figure 4:**
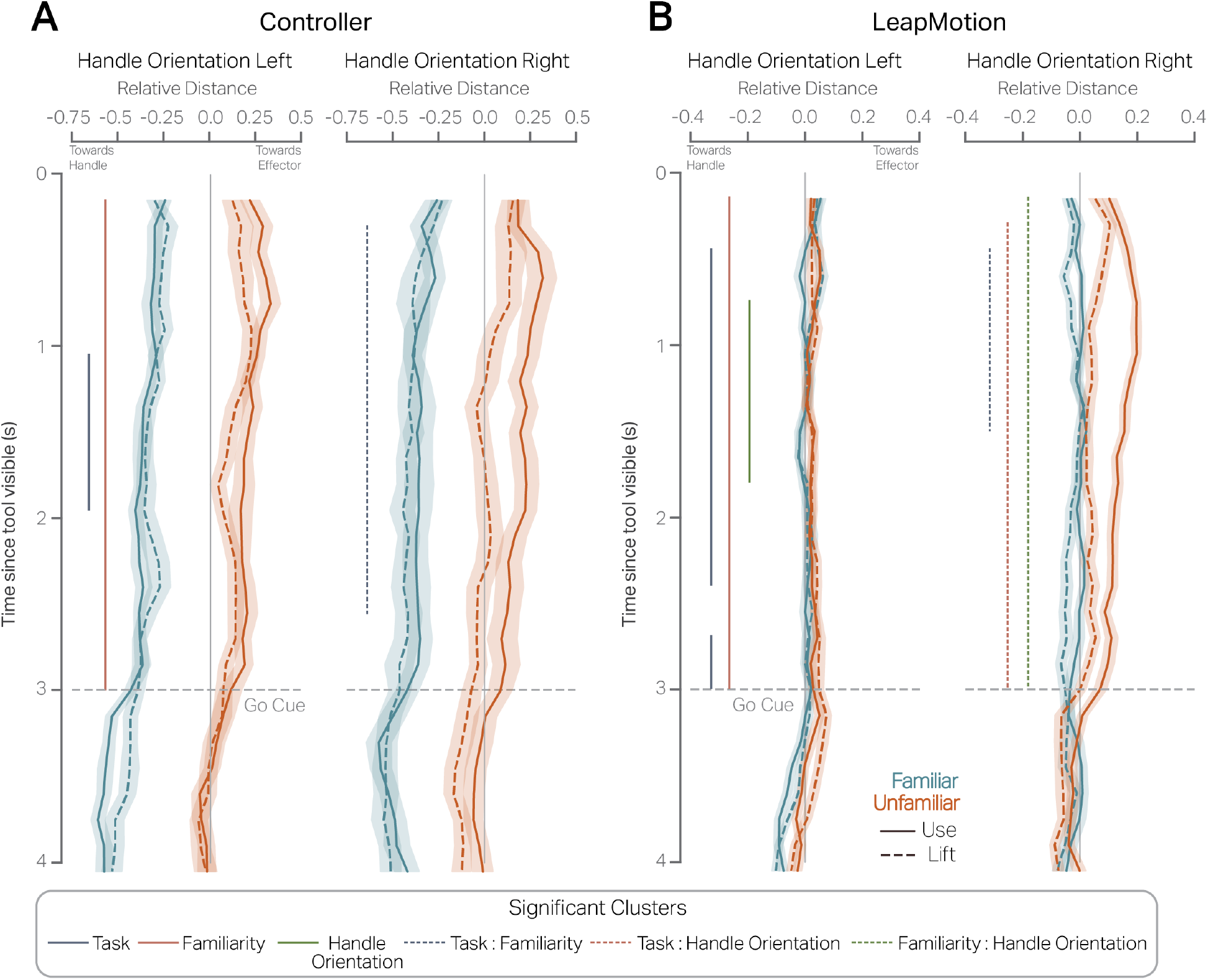
Eccentricity of fixations on the tool models. The negative values of the abscissa correspond to fixations towards the handle, the positive values refer to fixations towards the tool effector, and zero represents the center of the tool. The ordinate axis refers to the time elapsed since the tool is visible on the virtual table. The go cue is given to participants at 3s after which they can start interacting with the tool. The blue lines correspond to the FAMILIAR tool and red to the UNFAMILIAR tools. The error bars represent the standard error of the mean across subjects. The vertical solid lines correspond to the significant time clusters for main effects and the vertical dashed lines to the interactions. **Panel A** shows the findings from experiment-I and the two handle orientations. **Panel B** shows the findings from experiment-II and the two handle orientations.

Similarly, figure 4B shows the eccentricity of fixations from the center of the tool during the 3s period when the subjects studied the tool in experiment-II. The differences in task were significant in two time periods 0.45s to 2.4s, p-value< 0.001 and from 2.7s to 3s, p-value=0.03. The differences in familiarity were significant from 0.15s to 3s, p-value < 0.001. Furthermore, the differences in handle orientation were significant from 0.75s to 1.8s, p-value < 0.001. A significant interaction of task and familiarity from 0.45s to 1.5s with p-value <0.001. There were also significant clusters in the interaction of task and handle orientation, from 0.3s to 3s with p-value<0.001 and for tool familiarity and handle orientation from 0.15s to 3s with p-value < 0.001.

## 4 Discussion

The primary aim of this study is to investigate how gaze-based strategies vary for tasks, tool familiarity, and manual planning in naturalistic settings. With our study, we successfully added to the current body of research in two important ways. Firstly, irrespective of the realism of the action affordance in virtual environments, the number, and location of anticipatory fixations were modulated by goal-oriented factors of task and tool familiarity. Secondly, anticipatory fixations related to proximal manual planning were only seen when the setup allowed for more realistic action affordances with the virtual hand mimicking finer hand and finger movements. In sum, proximal and distal goal-oriented planning is highly contextualized to the realism of action/interaction afforded by the environment.

We conducted two experiments to disentangle the role of action affordance for goal-oriented planning. Participants interacted with 3D tool models using VR controllers in a low realism setup, which produced a virtual grasp by pulling their index fingers. Here, we showed that the odds of sampling visual information from the mechanical properties of a tool are different based on the specificity of the task. Moreover, given tool familiarity, the odds of fixating on the effector increased for unfamiliar tools. Tool-specific knowledge also played a major role when subjects were instructed to produce tool-specific movements. Moreover, the spatial orientation of the tool did not affect the odds of fixations for the tool effector. In sum, with the preparation of a symbolic grasping action, fixations were affected by distal goal-oriented factors of task and tool familiarity.

In a high realism setup, participants interacted with tool models by producing an actual grasp over the tools. The results were similar to the first experiment. However, we additionally found a significant effect of spatial orientation of the tool where the odds of fixations in favor of the tool effector decreased when the tool was presented incongruent to the subjects’ handedness. These results suggest that fixations are directed towards the handle of the tool in anticipation of planning the proximal goal of an optimal grasp. Interestingly, the optimal grasp planning is initiated from the beginning until the end of the viewing time window and might be more critical than inspecting the tool effector to produce the correct action. Taken together, the preparation of a realistic grasping action modulated anticipatory fixations related to both proximal and distal goal planning.

These results are in line with the findings reported by Belardinelli et al. (2016a). They investigated behavioral responses to task and tool-based affordances in a lab where subjects responded to stimuli images on a computer screen and pantomimed their manual actions. Moreover, they presented the tools with the handle always oriented on the right and congruent to the subject’s handedness. Our results suggest that well before action initiation, subjects had to substantially plan their hand movement on the tool to interact with it. This effect is indicative of an end-state comfort planning (Herbort and Butz, 2012) where both proximal and distal goal-oriented planning interacts to modulate anticipatory fixations. From the perspective of ecological validity, our findings give a fuller view of how different planning strategies are needed to produce relevant action. Our study shows that within a naturalistic setting, task, tool familiarity, and the spatial orientation of the tool affect the planning and production of relevant actions. Hence, our study offers a veridical and ecological valid context to aspects of anticipatory behavior control.

Studies in eye-hand coordination (Johansson et al., 2001; Lohmann et al., 2019; Belardinelli et al., 2018) have shown that eye movements are predictively made towards the grasp contact points. Furthermore, Flanagan et al. (2006) proposed that predictions are made in an event-oriented manner and are at the heart of successful control strategies for object manipulations. They posit that predicted sensory events are compared with actual events like grasping, lifting, moving the object to monitor task progression. In contrast, Iacoboni et al. (2005) and Wohlschläger et al. (2003) showed that goal-oriented planning is specified at an abstract level rather than at the movement level. Our results suggest that the anticipatory gaze behavior specific to task and tool familiarity is seen only when additional grasp control planning is not needed. Inversely, optimal motor control might supersede planning based on other distal goals. Here, we make the case that predictions are made for action outcomes at various scales, and that eye movements are used to plan both optimal grasp control and task-specific requirements well before action initiation.

Our study adds to the growing body of evidence that anticipation and prediction are at the core of cognition (Pezzulo et al., 2007). Motor theories of cognition have proposed that simulations of actions reuse internal models of motor commands to effect multiple predictions (Jeannerod, 2006). The simulation of action theory has been used to explain numerous phenomena of planning, prediction of external events, visual perception, and imitation. Hoffmann (2003) introduced anticipatory behavior control as the mechanism by which action-effect representations are activated by the need for an effect-related goal and contingent stimuli. Furthermore, Pezzulo et al. (2021) recently proposed that generative models provide top-down predictive signals for perception, cognition, and action during active tasks and these signals are otherwise weak and/or absent when the brain is at rest or the stimuli are weak. Our study shows that anticipatory behavior is tightly linked to the production of task-relevant actions and contextualized to the realism of the action affordance.

Notably, our study shows that different constraints on the method of interaction can also result in different anticipatory behavioral responses. From the perspective of Gibson (1977), the affordances of the environment are tightly linked to the actions that one can perform in it. Similarly, O’Regan and Noë (2001) posited that actions constitute the cognitive processes that govern relevant sensorimotor contingencies. In our study, the production of relevant actions significantly modulated the visual sampling of the tool parts in accordance to goal-oriented factors such as task and tool familiarity irrespective of the action affordance. Taken together, our study shows that some aspects of anticipatory gaze are dependent on the realism of the action afforded by the environment.

We conducted the present study in virtual reality, which is still a burgeoning technology for vision research. While VR environments pose an exciting avenue of research, there are still limitations that practitioners must face while conducting experiments in these scenarios. First, the naturalistic setting of both experiments I and II afforded natural head movements. To maintain optimal quality over the data, we asked the participants in the study to make limited head movements. Additionally, we presented the tools and the task cues not to cause extreme pitch head movements. Secondly, mobile eye-trackers can be error-prone and might suffer from variable sampling rates (Ehinger et al., 2019) or calibration errors due to slippage (Niehorster et al., 2020). To mitigate any calibration errors, we also made sure that we calibrated the eye-trackers at regular intervals. Thirdly, both controller-based and camera-based VR interaction methods are still new technology. It could have been challenging for participants to get used to, even though we made sure they practiced the interaction method before the experiment. While we simulated grasping the tool using LeapMotion’s gesture recognition and were able to produce a more realistic action affordance through mimicking finer hand and finger movements, it is still an inadequate substitute for a real grasp where the tactile feedback of the tool in hand might elicit more accurate responses. For example,Ozana et al. (2018) showed that grasping movements within a virtual environment differ both quantitatively and qualitatively from typical grasping. Lastly, there are obvious differences in the realism of the two virtual environments used in the study in terms of the visual scene. While there are visible differences between the environments, we see that there are no significant differences between the percentage of fixations allocated to the background vs. tool for both experimental settings. Hence, we contend that the differences in the eye movement behavior reported in the study are largely a consequence of the differences in the action affordance and much less because of mere visual differences. In light of these limitations, we know that our study must be considered from a nuanced perspective. Furthermore, there is still room for replicating our study with novel and more realistic interaction methods.

There are still some open questions pertaining to anticipatory behavior elicited by tool interactions. Firstly, while our study distinguishes between levels of action affordances, future work can look at goal-oriented planning for passive observers at both proximal and distal levels. Secondly, it would be interesting to dive deeper into the predictive brain signals that give rise to the present oculomotor behaviors. Our study provides a first step towards distinctly investigating proximal and distal goal-oriented planning.

## 5 Conclusion

The present study gives a veridical and ecologically valid context to planning and anticipatory behavior. Our results support the hypothesis that eye movements serve the cognitive function of actively sampling information from the environment to produce relevant actions. When semantic information about the object is not readily available, eye movements are used to seek information from its mechanical properties from specific locations. Furthermore, we show that fixations are made in a goal-oriented way in anticipation of the relevant action. When considering the realism of the action affordance, our results show that eye movements prioritize proximal goals of optimal grasp over task-based demands. Lastly, our study is at the frontiers of naturalistic vision research, where novel technologies can be harnessed to answer questions that were previously far-fetched.

## Author Contributions

AK, PK: conceived and designed the study. TS, PK: Procurement of funding. NG, AK, FNN: programmed the controller study. SB, AK, FNN: programmed the LeapMotion study. NG, SB: data collection. AK: data analysis. AK: initial draft of the manuscript. AK, SB, NG, FNN, TS, PK: revision and finalizing the manuscript. All authors contributed to the article and approved the submitted version.

## Acknowledgement

We are grateful for the financial support by: the German Federal Ministry of Education and Research for the project ErgoVR (Entwicklung eines Ergonomie-Analyse-Tools in der virtuellen Realität zur Planung von Arbeitsplätzen in der industriellen Fertigung)-16SV8052; German Research Foundation (Deutsche Forschungsgemeinschaft, DFG) project number GRK2340/1 (DFG Research Training Group Computational Cognition).

## Footnotes

1 https://enterprise.vive.com/us/product/vive-pro-eye-office/

2 Unity, www.unity.com

3 SRanipal, http://developer.vive.com/resources/vive-sense/sdk/vive-eye-tracking-sdk-sranipal/

4 SteamVR, https://valvesoftware.github.io/steamvr_unity_plugin/articles/Quickstart.html

5 LeapMotion Unity modules, https://developer.leapmotion.com/unity

## Notes

### Competing Interest Statement

The authors have declared no competing interest.

### Summary of Updates

Updated the title and the discussion section after internal reviews

